# A model for generating differences in microtubules between axonal branches depending on the distance from terminals

**DOI:** 10.1101/2022.06.13.496038

**Authors:** Chiaki Imanaka, Satoshi Simada, Shino Ito, Marina Kamada, Tokuichi Iguchi, Yoshiyuki Konishi

## Abstract

In the remodeling of axonal arbor, the growth and retraction of branches are differentially regulated within a single axon. Although cell-autonomously generated differences in microtubule (MT) turnover are thought to be involved in selective branch regulation, the cellular system whereby neurons generate differences of MTs between axonal branches has not been clarified. Because MT turnover tends to be slower in longer branches compared with neighboring shorter branches, feedback regulation depending on branch length is thought to be involved. In the present study, we generated a model of MT lifetime in axonal terminal branches by adapting a length-dependent model in which parameters for MT dynamics were constant in the arbor. The model predicted that differences in MT lifetime between neighboring branches could be generated depending on the distance from terminals. In addition, the following points were predicted. Firstly, destabilization of MTs throughout the arbor decreased the differences in MT lifetime between branches. Secondly, differences of MT lifetime existed even before MTs entered the branch point. In axonal MTs in primary neurons, treatment with a low concentration of nocodazole significantly decreased the differences of detyrosination (deTyr) and tyrosination (Tyr) of tubulins, indicators of MT turnover. Expansion microscopy of the axonal shaft before the branch point revealed differences in deTyr/Tyr modification on MTs. Our model recapitulates the differences in MT turnover between branches and provides a feedback mechanism for MT regulation that depends on the axonal arbor geometry.

## 1. Introduction

In the nervous system, axons elaborate branches to form arbors that innervate multiple target cells (Gibson and Ma, 2011; Kalil and Dent, 2014). Remodeling of axonal arbor has been observed in multiple brain regions even in the adult stage (Stettler et al., 2006; Nishiyama et al., 2007), indicating it plays critical roles not only in the establishment of neuronal wiring but also for neuronal plasticity. This process involves the formation and extension of branches in which some branches are simultaneously eliminated or shortened by retraction. The mechanisms whereby neurons selectively regulate growth or retraction remain to be elucidated.

Cell extrinsic cues from surrounding tissues or target cells play roles in selective branch regulation. In the development of subcortical projection, layer 5 pyramidal neurons extend collateral branches to multiple brain areas. The extension and elimination of specific branches establish neuronal subtype specific axonal branch patterns (O’Leary and Koester, 1993). It is considered that this process is regulated by axon guidance molecules expressed in a region-dependent manner (Iguchi et al., 2021). In the case of motor neuron axons innervating the muscles, multiple axons contact the same synaptic site on muscles by forming highly branched terminal arbors at the neonatal stage. During postnatal development, many branches are retracted while the remaining branches are simultaneously stabilized so that each muscle fiber makes a synapse with a single axon (Walsh and Lichtman, 2003). Blocking the activity of postsynaptic muscle fibers prevents this process. A recent study revealed that MTs in retracting branches are selectively severed by spastin, and synaptic elimination and retraction of branches is not blocked but significantly delayed in spastin knockout mice (Brill et al., 2016). These observations suggest that signals from the neuromuscular junction may trigger the elimination of branches by destabilizing MTs.

In addition to extracellular signals, cell-autonomous systems are involved in the selective branch regulation in axons. At neuromuscular junctions, a motor neuron eliminates branches from some muscular sites while simultaneously stabilizing the remaining branches (Lichtman and Colman, 2000). It has been observed that isolated neurons in culture differentially regulate the growth and retraction of branches in the same axon in the absence of extrinsic cues (Ruthel and Hollenbeck 2000, 2003). Furthermore, growth competition between neighboring branches had been observed (Hutchins and Kalil, 2008). Although a difference in frequency of localized spontaneous Ca^2+^ transients is suggested to mediate competitive signals between branches (Hutchins and Kalil, 2008), the mechanism by which neurons select particular branches remains unknown.

During the elongation of MTs, α/β-tubulin heterodimers are incorporated into the plus end as a GTP-bound form. After incorporation, GTP bound to β-tubulin is stochastically converted to GDP. Exposure of GDP-tubulins destabilizes the plus end of MTs and causes shrinkage, which is rescued by incorporation of new GTP-tubulins, a behavior known as dynamic instability (Mitchison and Kirschner, 1984). In addition, tubulins receive post-translational modifications that depend on their GTP/GDP state and affect the functions of kinesins and MAPs (Gundersen et al., 1987; Janke and Magiera, 2020). We have reported that the motor domain of MT motor kinesin-1 accumulates in a subset of arbor branch terminals and that contributes to inhibiting the retraction of branch terminals. In neighboring branch terminals, kinesin-1 is preferentially transported in longer branches than shorter branches (Seno et al., 2016). We also found that longer branches contain more stable MTs, which are reported to enhance kinesin-1 mediated transport. These observations indicate that there is a positive feedback loop between MT stability and branch length. However, the mechanism by which branch dependent differences of axonal MTs are generated has not been elucidated. In this study, we aimed to generate a model for a cell-autonomous system for creating branch dependent differences of MTs.

## 2. Results

### 2.1 Differences of MT lifetimes between branches can be generated by a simple model with MT dynamic instability

We modeled terminal axonal arbor with longer and shorter branches of length *L*_*l*_, and *L*_*s*_ (Fig. 1A, B). Differences of MT stability between branches can be caused by region dependent regulation (Fig. 1A). This system requires spatial cues to assign specific branches. According to the cell length dependent model (Seetapun and Odde, 2010), as the process becomes longer, stable microtubules are accumulated because of a greater traveling time from the soma to the terminal, where most MTs undergo catastrophe. To test whether the distance from terminals can generate branch dependent differences of MT lifetime, we adopted this model in axonal arbors. Because MTs in mature axons are segmented (Yu and Baas, 1994; Kapitein and Hoogenraad, 2015) and their distribution regulation is unknown, we assumed that MT minus-ends were randomly distributed in our model arbor. MTs repeatedly grow and shorten by dynamic instability, and at the axonal terminal, MTs undergo shrinkage by catastrophe (Seetapun and Odde, 2010). In our model, dynamic instability and continuous catastrophe at terminals would generate a gradient of MT lifetimes along axons, and branch dependent differences could be generated according to the distance from terminals (Fig. 1B).

**Fig. 1.**
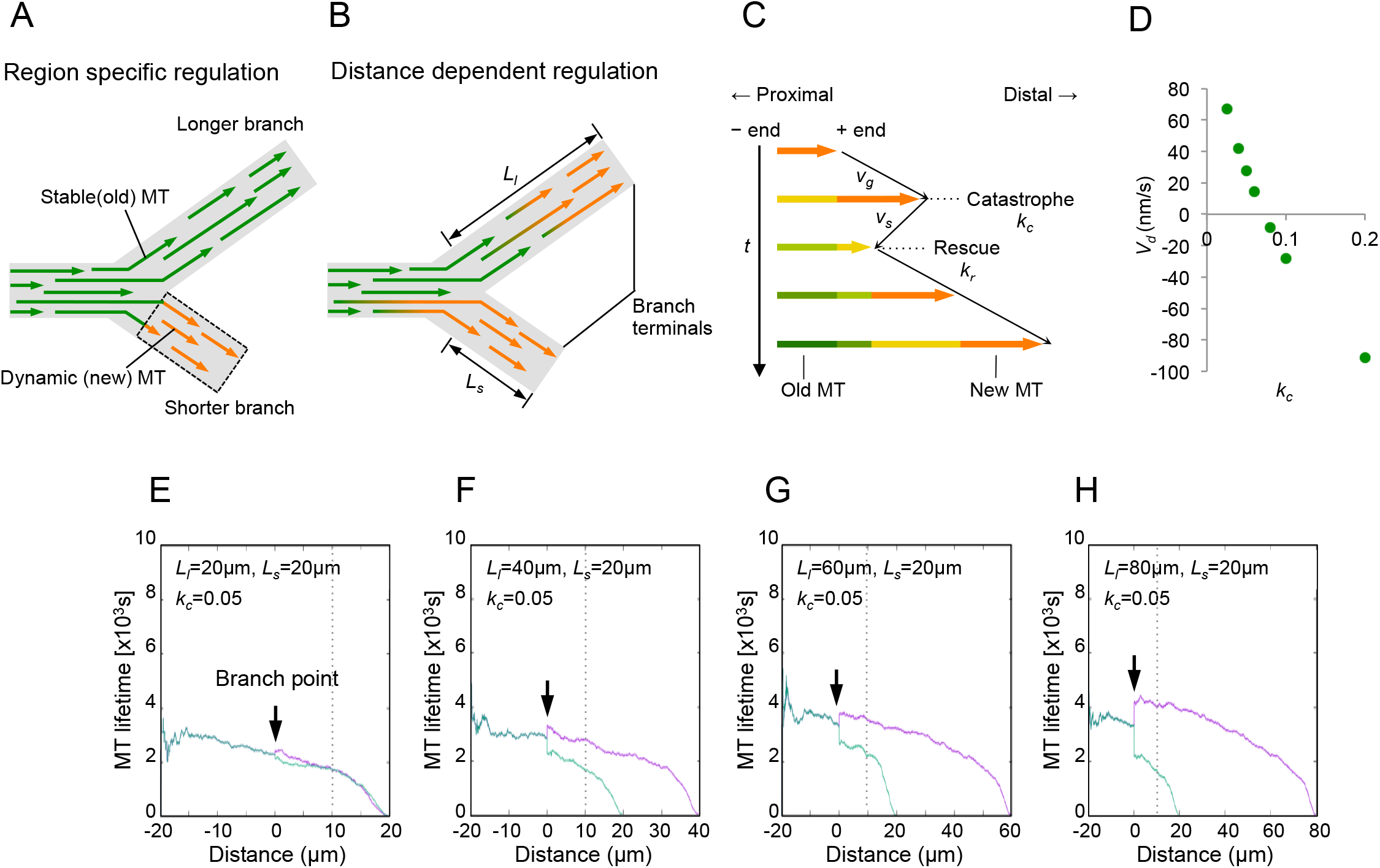
Models for the generation of branch-dependent differences of MTs in axonal arbors. (A, B) Adjacent branches with different lengths were assumed in the model. Longer and shorter branches tend to be enriched with stable MTs and unstable MTs, respectively. This difference can be explained by region specific regulation (dashed rectangle) of MT stability in a particular branch (A) or regulation depending on the distance from axonal terminal ends (B). *L*_*l*_ and *L*_*s*_ represent the length of longer and shorter branches, respectively. (C) A schematic diagram of dynamic instability and lifetime of MTs. Each MT stochastically switched between growth (at velocity *v*_*g*_) and shrinkage (at velocity *v*_*s*_) phases depending on the frequency of catastrophe (*k*_*c*_) and rescue (*k*_*r*_), generating a gradient from old to new. (D) A plot of the mean change in MT length (*V*_*d*_) versus the corresponding values of *k*_*c*_ (E–H). The effect of branch length (*L*_*s*_ = 20 µm, *L*_*l*_ = 20, 40, 60, 80 µm) on the MT lifetime in models with distance dependent regulation at *k*_*c*_ = 0.05 and *k*_*r*_ = 0.15. The MT lifetime in each position relative to the branch point (arrow) was calculated. The density of MT minus-ends in each branch was set to 100/10 µm. Values calculated from longer branches and shorter branches were colored by magenta and light green, respectively. The dotted lines indicate the positions 10 µm distal to the branch point.

Each MT stochastically switches between states of growth or shrinkage (Janulevicius et al., 2006; Seetapun and Odde, 2010), depending on the frequency of rescue (*k*_*r*_) and catastrophe (*k*_*c*_), respectively (Fig 1C). Consequent to the dynamic instability of each MT, a difference of lifetime is created not only between MTs but also within a single MT (Fig 1C). In this study, we monitored the existence time of each MT segment. Velocity of MT shrinkage (*v*_*s*_) and *k*_*r*_ were set as in the previous study (Seetapun and Odde, 2010) (Table 1). For the velocity of MT growth (*v*_*g*_), we used the value obtained by imaging of plus ends of growing MTs by using end-binding 3 (EB3) in branched axonal arbor (Seno et al., 2016), which was slightly lower than the value obtained from newly generated axons (120 nm/s vs. 150 nm/s). For *k*_*c*_, the mean value calculated from the data of our previous study was much lower than that from the study of newly generated axons (0.03 s^−1^ vs. 0.1 s^−1^) (Seetapun and Odde, 2010; Seno et al., 2016). This difference may have arisen from differences in axonal maturity and other situations. The mean change in MT length (*V*_*d*_) can be described by the following equation (Janulevicius et al., 2006; Seetapun and Odde, 2010):

**Table 1.**

| Constant | Definition | Value |
| --- | --- | --- |
| $v_g$ | Velocity of MT growth | 120 nm/s |
| $v_s$ | Velocity of MT shrinkage | 250 nm/s |
| $k_c$ | Rate of MT catastrophe | $0.05\text{--}0.2 \text{ s}^{-1}$ |
| $k_r$ | Rate of MT rescue | $0.15 \text{ s}^{-1}$ |

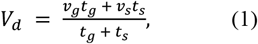

where *t*_*g*_ = 1/*k*_*c*_ is the growth time, and *t*_*s*_ = 1/*k*_*r*_ is the shrinkage time. By increasing *k*_*c*_, the net change of MT length was altered from growth (*V*_*d*_ > 0) to shortening (*V*_*d*_ < 0) (Fig. 1D). From the equation, increasing the *k*_*r*_ would have the opposite effect on *V*_*d*_. We treated *k*_*c*_ as an adjustable parameter by fixing other parameters to investigate the effect of MT dynamics on branch-dependent differences.

Using the branch model with *k*_*c*_ = 0.05 s^−1^, such that *V*_*d*_ = 27.5 nm/s, the model predicted a branch dependent difference of MT lifetime (i.e., mean existence time). Here, *L*_*s*_ was set to 20 µm, and *L*_*l*_ was variable. To obtain an average, 100 MTs per 10 µm in both branches were assumed. A difference between the shorter branch and longer branch was detected when *L*_*l*_ = 40 µm, and the difference increased as *L*_*l*_ became longer (Fig 1E–H). The MT lifetime gradually decreased towards the ends with a similar slope for both branches, as experimentally observed (Seno et al., 2016). The MT lifetimes calculated from experimental data (10 µm distal to the branch point) were 3.5×10^3^ min and 1.1×10^3^ min for longer and shorter branches (Seno et al., 2016), which were not inconsistent with the values of MT age observed in the model (Fig 1E–H). These results suggest that the axonal arbor model based on MT regulation according to the distance from terminals recapitulates the differences of MT aging between neighboring branches.

### 2.2 Differences in MT lifetimes between branches can be reduced by MT destabilization in the model

Next, we simulated MT lifetime with *k*_*c*_ = 0.1 s^−1^, such that *V*_*d*_ = −28 nm/s, and other parameters fixed as in Figure 1E–H. We found that at the base of the branches (10 µm distal to the branch point), the differences between longer and shorter branches became smaller if anything (Fig. 2A–D). At *k*_*c*_ = 0.2 s^−1^ (*V*_*d*_ = −98.4 nm/s), MT lifetime took a very small value with high variation, throughout the arbor, reflecting that only a small portion of MTs remained (Fig. 2E–H). Figure 2I shows a profile of the differences in MT lifetime between branches with different collapse frequencies (*k*_*c*_ = 0.05 s^−1^ and 0.1 s^−1^) and branch lengths (*L*_*l*_ = 20–100 µm, *L*_*s*_ = 20 µm) obtained from three simulations. At *k*_*c*_ = 0.05 s^−1^, the differences between branches appeared at *L*_*l*_ = 30µm and *L*_*s*_ = 20 µm (ratio of MT-lifetime of shorter/longer = 0.74 ± 0.02). When the length of longer branches was increased, the ratio (shorter/longer branches) of MT lifetimes became lower (at *L*_*l*_ = 100 µm, shorter/longer = 0.41 ± 0.04). The ratios of MT lifetimes were always less than 1 using the model, in which we set the density of MTs to be 100 MT/10 µm. However, only several MTs are expected to exist in the 10 µm segment of mature axons (Yu and Baas, 1994; see below). We, thus, performed a simulation to lower the MT density from 100/10 µm to 5/10 µm (Fig. 2J). Although the ratios of MT lifetimes were less than 1 in most cases, reversal of MT lifetimes between short and long branches was observed in some cases with 5 MTs/10 µm. This indicates that the difference between long and short branches was not constantly maintained in this model, but was stochastically switchable, as observed in actual axons. At *k*_*c*_ = 0.1 s^−1^, the ratio of MT lifetimes was lower (at *L*_*l*_ = 100 µm, shorter/longer = 0.77 ± 0.05) and more frequently reversed between short and long branches when MT number was reduced (Fig. 2J, K). Collectively, in the current model arbor, increasing the catastrophe of MT reduced the difference in MT lifetimes between shorter and longer branches.

**Fig. 2.**
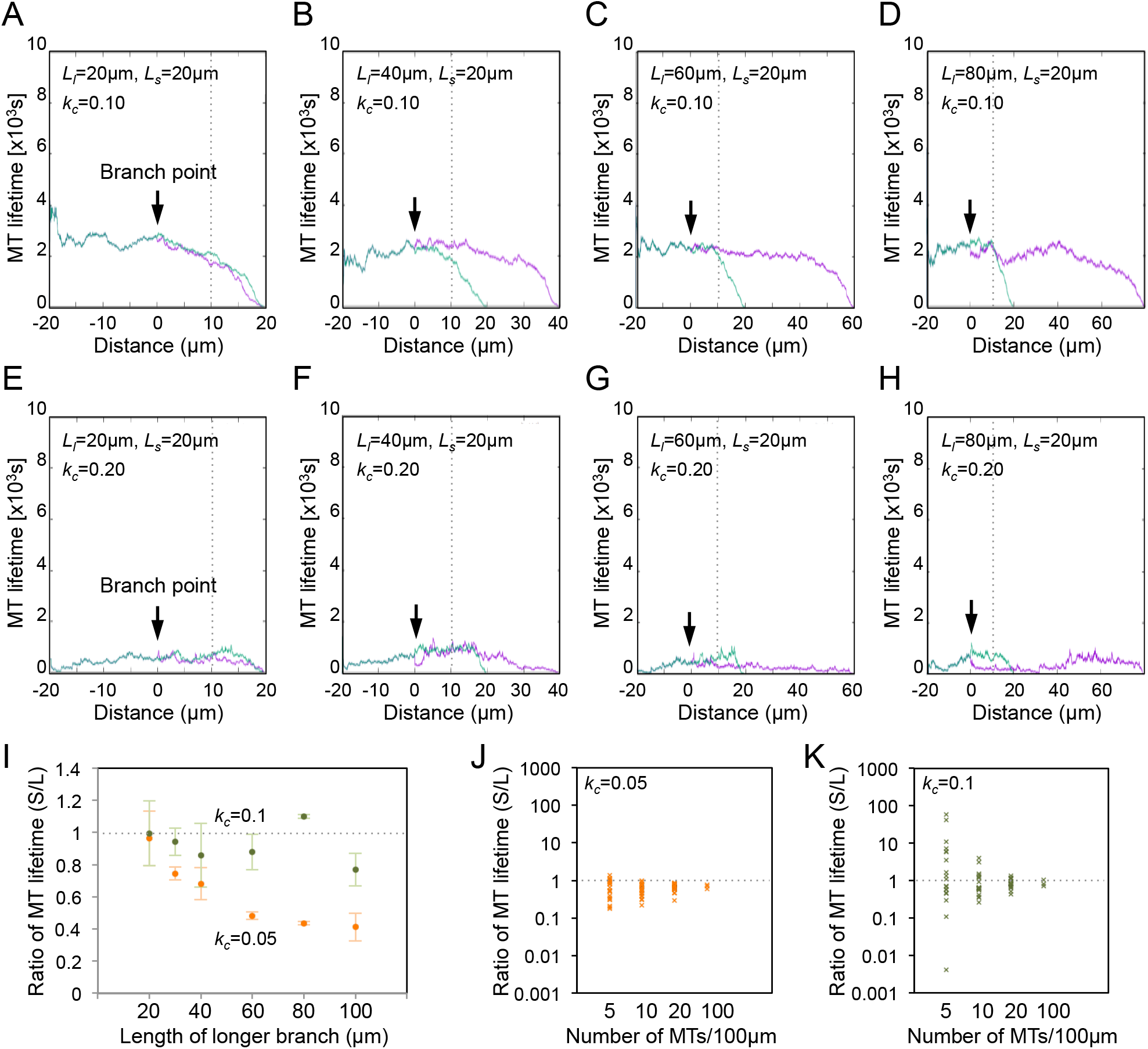
Effect of *K*_*c*_ on MT differences between branches of the axonal model. (A–D) Effect of branch length (*L*_*s*_ = 20 µm, *L*_*l*_ = 20, 40, 60, 80 µm) on MT lifetime in the models at *k*_*c*_ = 0.1. Differences between longer and shorter branches were smaller compared to the case of *k*_*c*_ = 0.05. (E–H) Simulated relative MT lifetimes at *k*_*c*_ = 0.2 revealed low values in both shorter and longer branches. (I) A profile of simulated MT lifetimes at *k*_*c*_ = 0.05 (orange) and *k*_*c*_ = 0.1 (green) in the model axonal arbor (*L*_*s*_ = 20 µm, *L*_*l*_ = 40 µm). Values were obtained from *n* = 3 simulations and are represented as the mean ± 95% confidence interval (CI). (J, K) Simulated relative MT lifetimes in the model (*L*_*s*_ = 20 µm, *L*_*l*_ = 40 µm) at *k*_*c*_ = 0.05 (J) and *k*_*c*_ = 0.1 (K) in which 5 to 100/10 µm MT minus-ends were placed in both branches. Cross marks represent the value obtained from one simulation. A reduction of the number of MTs increased the number of cases in which the lifetimes of MTs reversed between short and long branches.

### 2.3 MT destabilization with nocodazole treatment reduces the difference in MT lifetimes between branches

To determine whether MT destabilization reduces the difference in MT lifetimes between longer and shorter branches in actual axonal arbor as in the model, we investigated MTs states in axons of cultured neurons. Cerebellar granule neurons (CGNs) were cultured at low density to avoid intercellular interactions (Kubota et al., 2013), and neurons were treated with nocodazole to cause destabilization of MTs at 250 nM because high concentrations of nocodazole cause beading and degradation of axons (Datar et al., 2019). In the dimer state, α-tubulin receives a tyrosine residue at the carboxy terminal to become tyrosinated (Tyr) by the action of tubulin of tyrosine ligase (Ttl) (Ersfeld et al., 1993; Erck et al., 2005), whereas in the polymerized state, α-tubulin is detyrosinated (deTyr) by removal of the tyrosine residue by vasohibins or microtubule associated tyrosine carboxypeptidase (Aillaud et al., 2017, Nieuwenhuis et al., 2017; Landskron 2022). Whereas newly generated MTs are enriched with Tyr-tubulin, the deTyr/Tyr ratio increases with the lifetime of MTs. Thus, deTyr/Tyr ratio of α-tubulin has been widely used as an indicator for the turnover of MTs (Kreis et al., 1987; Brown et al., 1993; Hammond et al., 2010; Moutin et al., 2021). CGNs (4 days *in vitro*) that had been treated with nocodazole for 2.5 h were fixed and stained using an antibody against Tyr- and deTyr-tubulins. We observed arbor terminals that had two neighboring branches of different length. In control neurons, the deTyr/Tyr ratio of MTs was high at axonal terminals and gradually increased in a distance-dependent manner. The ratio was greater in longer branches at the same distance from a branch point, as reported previously (Seno et al., 2016) (Fig. 3A). In nocodazole treated neurons, the deTyr/Tyr ratio tended to be low even in longer branches (Fig. 3A). We quantified the deTyr/Tyr ratio of branch pairs of which the length of shorter branches was more than 10 µm and the length of longer branches was more than twice as long as the shorter ones (Fig. 3B, C). At the base of branches (up to 10 µm from the branch point), longer branched exhibited significantly higher deTyr/Tyr ratios compared with shorter branches, but nocodazole treatment reduced the differences between longer and shorter branches in control neurons (median 1.2 vs. 0.7, p < 0.001, in control (DMSO); median 1.0 vs. 0.8, p > 0.05 in 250 nM nocodazole, Wilcoxon signed-rank test). These experimental results indicate that destabilization of MTs in actual axonal arbor reduces the differences in MT lifetimes between shorter and longer branches, as shown in the model.

**Fig. 3.**
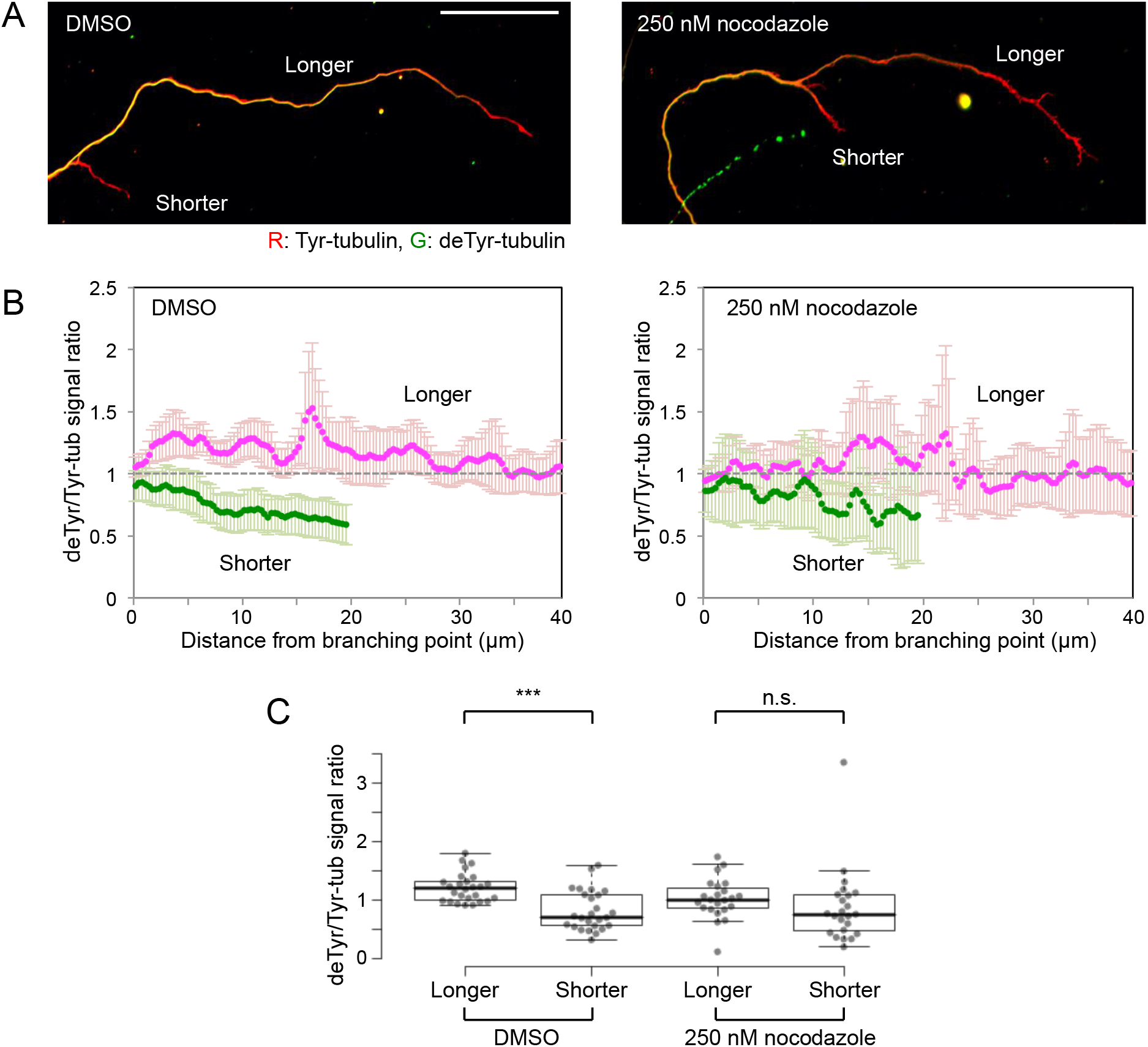
Nocodazole treatment reduced the differences of deTyr-/Tyr-tubulin ratios between branches of axonal arbors. (A) Axons of CGNs that had been treated with DMSO or 250 nM nocodazole were subjected to immunocytochemistry using antibodies for deTyr- and Tyr-tubulins, a marker for old or new MTs, respectively. The scale bar indicates 50 µm. (B) Quantification of the deTyt-/Tyr-tubulin ratio in shorter (light green) and longer (magenta) branches. The values in each region were normalized to the signal at the branch point and are indicated as the mean ± 95% CI (*n* = 26 branch pairs in DMSO and *n* = 23 branch pairs in 250 nM nocodazole collected from three independent experiments). (C) Mean values up to 10 µm from branch point in (B) were compared. deTyt-/Tyr-tubulin ratio was significantly higher in longer branches (p < 0.001), whereas the nocodazole treatment reduced the difference (p > 0.05). The whiskers show the smallest and largest values within a distance of 1.5 times the interquartile range above and below the limits of the box. Data points are represented by dots.

### 2.4 There are differences in deTyr/Tyr between MTs that enter longer and shorter branches even before the branch point

In the model in which MTs are regulated depending on the distance from arbor terminals, we noticed that differences of MT lifetimes between longer and shorter branches are generated even at the branch point (Fig. 1E–H). In the shaft before the branch point, because the lifetimes of MTs that enter the two branches were averaged, a difference of MT age could exist before the segregation into the two branches (Fig. 1B). We investigated in actual axonal arbors whether there was a difference between MTs that entered into long and short branches at the axonal shaft before the branch point. In normal microscopy, although differences between longer and shorter branches can be detected by proximity to the branch point, the deTyr/Tyr of each MTs cannot be detected because of the limitation of spatial resolution (Fig. 3A, B). Thus, we performed expansion microscopy (ExM) (Chen et al., 2015) to observe the deTyr/Tyr of MTs around the branch point. Figure 4 is an example of axonal arbor that has longer and shorter branches. Several MTs with different deTyr/Tyr ratios were detected in longer and shorter branches, and MTs in the shorter branch exhibited lower deTyr/Tyr ratios (position (a) of Fig. 4A, B). At the axonal shaft before the branch point, although overall deTyr/Tyr ratios tended to be higher than in the branches, differences in MT modification could already be detected before MTs entered the branch point (position (b) of Fig. 4A, B). This observation is consistent with the simulation results (Fig. 1E–H); thus, in axonal branches of isolated neurons *in vitro*, the lifetimes of MTs are regulated according to the distance from terminals rather than the region dependent regulation (Fig. 1A, B).

**Fig. 4.**
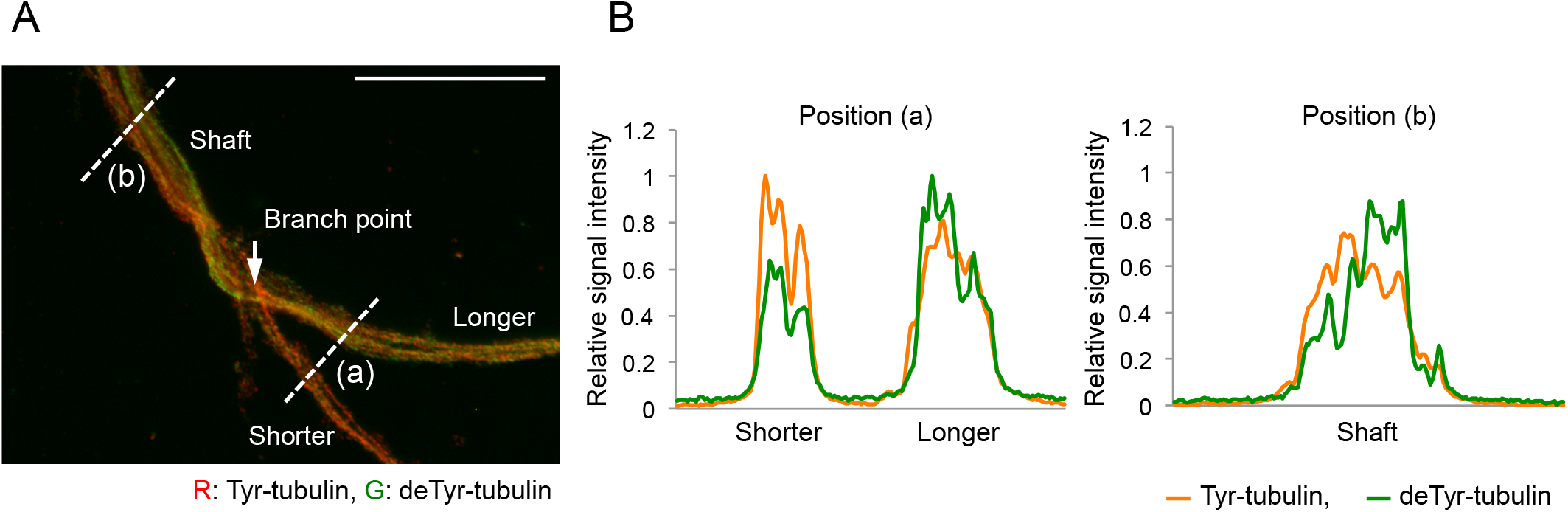
Differences of MTs entering adjacent branches existed even before the branch point. (A) ExM of deTyr-(green) and Tyr-(red) tubulins in an axonal arbor that had longer and shorter branches. An image of around the branch point (arrow) is shown. The scale bar indicates 10 µm. (B) deTyt- and Tyr-tubulin signals of MTs in branches (a) and before the branch point (b) in (A) are shown. The ratio of deTyt-/Tyr-tubulin varied between MTs, and there was a significant difference between the branches. Note that the segregation of MTs can be found before the branch point.

## 3. Discussion

Previous studies have revealed region dependent differences in MT turnover within a single neuron. In general, MTs in the shaft of axons tend to be more stable than those in axonal terminal and somatodendritic regions (Witte et al., 2008; Hammond et al., 2010). Within a single axonal arbor, MT turnover tends to be slower in longer branches compared with shorter branches (Seno et al., 2016). However, the feedback mechanism by which MT turnover is altered according to branch length has not been clarified. During the establishment of neuronal polarity, as a future axon extends beyond other minor processes, stable MTs accumulate in the axonal process. In this study, we adopted the cell length model (Seetapun and Odde, 2010) for microtubules in axonal branch arbors. The states of MTs in the model and actual axonal arbors were consistent in several aspects. The velocity and time of MT growth in the branch were the same within the axonal arbor. The age of MT gradually increased from the axonal terminal towards the branch point with a similar slope for both branches (Fig. 1E–H), as experimentally observed using Tyr- and deTyr-tubulin signals (Seno et al., 2016). Finally, a difference in the lifetimes of MTs between the longer and shorter branches was generated, and MT destabilization resulted in a reduction of the differences (Fig. 2I), as observed in cultured neurons (Fig. 3B, C). Because MT growth and stability may vary depending on the context, the current model may provide new perspectives in explaining differences in arbor morphology depending on neuronal subtypes or timing of development.

In polarized transport between axons and dendrites, besides on the differences in microtubules, axon initial segments, which function as gatekeepers, play pivotal roles (Konishi and Setou, 2009; Lewis et al., 2009; Song et al., 2009; Nakata et al., 2011). However, a less clear structural difference between axonal branches exists, presumably because the amounts of molecules to be transported rather than the types need to be regulated. In order to conduct differential sorting in axonal arbors, anterograde motors need to select a particular branch on which to transport. In our model, microtubule differences before the branch point may allow efficient selection of tracks to transport molecules to specific branches. Previous studies reported that kinesin-1 prefers MTs that contain detyrosinated and acetylated α-tubulins, indicators for stable MTs in cells (Reed et al., 2006; Dunn et al., 2008; Konishi and Setou, 2009). Recent studies also reported that MAP7 contributes to recruiting kinesin-1 on MTs at axonal branches (Tymanskyj et al., 2018). These factors may be orchestrated to achieve the traffic control within axonal arbors. Our present study provided a model that explains the mechanisms for the generation of branch dependent MT differences. Together with previous reports, the model provides insights into the cellular systems that control the traffic of molecules and the morphological changes in a branch dependent manner in axonal arbors.

## 4. Experimental procedure

### 4.1 Simulation of MT dynamics in axonal arbors

A program was written in the C language. The dynamic instability of MTs was simulated as described previously (Sprague et al., 2003; Seetapun and Odde, 2010) but with modifications. Because the MT minus-ends are sparsely distributed with an uncertain pattern in the axon of mature neurons, they were randomly distributed in the arbor. Each MT switches between growth and shortening states that were assumed to occur stochastically at frequency *k*_*c*_ (catastrophe: growth to shortening) and *k*_*r*_ (rescue: shortening to growth). In the growth state, MTs were allowed to elongate toward axonal terminals from the minus ends at velocity *v*_*g*_, whereas in the shortening state MTs shrank at *v*_*s*_ until the plus end reached the minus end. When the plus end reached the end of the axonal terminals away from the branch point with length *L*_*l*_ (longer branch) or *L*_*s*_ (shorter branch), MTs were immediately switched to the catastrophe state. During the simulation with 1 nm spatial steps and 1 s time steps, the existence time of each MT segment was recorded. After 10,000 cycles, the MT lifetime (mean time of all MTs at each position) was calculated.

### 4.2 Preparations of low-density culture neurons

CGNs were cultured at low density as described previously (Kubota et al., 2013). Mice were treated according to the institutional ethical guidelines of the University of Fukui (number: R02901). Postnatal day (P) 5–6 mice were obtained by crossing Jcl:ICR mice (CLEA Japan, Tokyo, Japan) that had been maintained at 18–28°C under a controlled light cycle (10 h light/14 h dark). Mice were decapitated, and isolated cerebella were digested with trypsin (TRL, Worthington, Lakewood, NJ). CGNs were suspended in Minimal Essential Medium (11090-081, Thermo Fisher Scientific, Waltham, MA) supplemented with 10% calf serum (SH30072.03, Thermo Fisher Scientific), penicillin (100 units/ml, P7794, Sigma-Aldrich, St. Louis, MO), streptomycin (0.1 mg/ml, P9137, Sigma-Aldrich), glutamine (2 mM, G8540, Sigma-Aldrich), and KCl (25 mM). CGNs were plated onto round cover glass (13 mm) that has been attached with paraffin dots as spacers and pre-coated with 15 µg/ml poly-L-ornithine at 2–4 × 10^4^ cells/well of a 24 well plate. After several hours of incubation (37°C, 5% CO_2_), CGNs on the cover slip were placed in another well containing a high density of CGNs (2.5 × 10^5^ cells/well) so that cells faced each other across the spacers to support survival (Kubota et al., 2013). Cytosine β-D-arabinofuranoside (10 µM, 034– 11,954, Sigma-Aldrich) was added to the culture at 1 day *in vitro*.

### 4.3 Immunocytochemistry

Immunocytochemistry of Tyr- and deTyr-tubulins was performed as described previously (Seno et al., 2016; Inami et al., 2018). CGNs were fixed for 20 min at room temperature with 4% paraformaldehyde (PFA, 163–18435, FUJIFILM Wako Pure Chemical Corporation, Osaka, Japan) in phosphate buffered saline (PBS). After fixation, CGNs were treated with 0.4% polyoxyethylene octylphenyl ether (169–21105, FUJIFILM Wako Pure Chemical Corporation) in PBS for 15 min at room temperature, followed by incubation for 1 h in blocking solution containing 5% goat serum (16210–064, Thermo Fisher Scientific), 3% bovine serum albumin (BAC62, Equitech-Bio, Kerrville, TX), and 0.02% polyoxyethylene sorbitan monolaurate (166–21115, FUJIFILM Wako Pure Chemical Corporation) in PBS. CGNs were then incubated with primary antibodies in blocking solution at 4°C overnight, followed by incubation with secondary antibodies for 2 h at room temperature. For primary antibodies, a monoclonal antibody against tyrosinated tubulin (1:2000; TUB-1A2, T9028, Sigma-Aldrich) and a polyclonal antibody against detyrosinated tubulin (1:1000; AB3201, Merck Millipore, Darmstadt, Germany) were used. For secondary antibodies, goat anti-mouse-IgG conjugated to Alexa Fluor 568 (1:1000; ab175473, Abcam, Cambridge, UK) and goat anti-rabbit-IgG conjugated to Alexa Fluor 488 (1:1000; A11008, Thermo Fisher Scientific) were used.

### 4.4 Expansion microscopy

Expansion microscopy (ExM) was performed as described previously (Chen et al., 2015, Chozinski et al., 2016). CGNs on the cover slip were fixed with a solution containing 3.2% PFA and 0.1% glutaraldehyde (072-02262, FUJIFILM Wako Pure Chemical Corporation) in PEM (0.1 M PIPES, pH 7.0, 1 mM EDTA, 1 mM MgCl_2_) for 10 min at room temperature. Following the immunocytochemistry as described above, CGNs were treated with 0.25% glutaraldehyde in PBS for 10 min and then washed three times with PBS. For gelation, 70 µl of monomer solution (2 M NaCl, 2.5% acrylamide, 0.15% N,N′-methylenebisacrylamide (011-08015 and 134-02352, FUJIFILM Wako Pure Chemical Corporation), 8.75% sodium acrylate (408220, Sigma-Aldrich) in PBS) added with 0.02% of tetramethylethylenediamine and ammonium persulfate (205-06313 and 012-08023, FUJIFILM Wako Pure Chemical Corporation) was applied on an O-ring (10 mm diameter, 1 mm thickness) placed on a piece of parafilm M (Amcor, Zurich, Switzerland). The cover slip was placed on the top with the cells facing down and incubated for 30 min at room temperature. The gels were digested with 8 units/ml of Proteinase K (075-02431, FUJIFILM Wako Pure Chemical Corporation) at 37°C for 1 h in digestion buffer (tris-acetate-EDTA buffer containing 0.5% polyoxyethylene octylphenyl ether and 0.8 M guanidine HCl (168-11805 and 075-02431, FUJIFILM Wako Pure Chemical Corporation)) and were placed in an excess amount of water for 2.5 h. The water was exchanged every 30 min.

### 4.5 Image processing and data analysis

Cell images were acquired using an Axiovert 200 M equipped with an MRm monochromatic digital camera (Carl Zeiss, Oberkochen, Germany) with 1388 × 1040 pixels using a 40× objective lens. Image data were processed by ImageJ software (National Institute of Health, Bethesda, MD). To quantify the signals of deTyr and Tyr tubulins, images of axonal arbor terminals, of which branches were more than 10 µm in length, were obtained, and the signal value along the axonal branch was obtained by plot profile. Image acquisition and data analysis were conducted in a blinded manner. For statistics, Wilcoxon rank sum test (R software) was used. No calculation for sample size predetermination was performed, and no data points were eliminated. No test for outliers was performed.

## Acknowledgements

This work was supported by JSPS KAKENHI Grant Numbers 17K07107 and 20K06889 (Y.K.).

## Abbreviations

MT: microtubule
CGN: cerebellar granule neuron
Tyr: tyrosination
deTyr: detyrosination
PBS: phosphate buffered saline
PFA: paraformaldehyde
DMSO: dimethyl sulfoxide
CI: confidence interval

